# MATEdb, a data repository of high-quality metazoan transcriptome assemblies to accelerate phylogenomic studies

**DOI:** 10.1101/2022.07.18.500182

**Authors:** Fernández Rosa, Tonzo Vanina, Simón Guerrero Carolina, Lozano-Fernandez Jesus, Martínez-Redondo Gemma I., Balart-García Pau, Aristide Leandro, Eleftheriadi Klara, Vargas-Chávez Carlos

## Abstract

With the advent of high throughput sequencing, the amount of genomic data available for animals (Metazoa) species has bloomed over the last decade, especially from transcriptomes due to lower sequencing costs and easier assembling process compared to genomes. Transcriptomic data sets have proven useful for phylogenomic studies, such as inference of phylogenetic interrelationships (e.g., species tree reconstruction) and comparative genomics analyses (e.g., gene repertoire evolutionary dynamics). However, these data sets are often analyzed following different analytical pipelines, particularly including different software versions, leading to potential methodological biases when analyzed jointly in a comparative framework. Moreover, these analyses are computationally expensive and not affordable for a large part of the scientific community. More importantly, assembled transcriptomes are usually not deposited in public databases. Furthermore, the quality of these data sets is hardly ever taken into consideration, potentially impacting subsequent analyses such as orthology and phylogenetic or gene repertoire evolution inference. To alleviate these issues, we present Metazoan Assemblies from Transcriptomic Ensembles (MATEdb), a curated database of 335 high-quality transcriptome assemblies from different animal phyla analyzed following the same pipeline. The repository is composed, for each species, of (1) a de novo transcriptome assembly, (2) its candidate coding regions within transcripts (both at the level of nucleotide and amino acid sequences), (3) the coding regions filtered using their contamination profile (i.e., only metazoan content), (4) the longest isoform of the amino acid candidate coding regions, (5) the gene content completeness score as assessed against the BUSCO database, and (6) an orthology-based gene annotation. We complement the repository with gene annotations from high-quality genomes, which are often not straightforward to obtain from individual sequencing projects, totalling 423 high-quality genomic and transcriptomic data sets. We invite the community to provide suggestions for new data sets and new annotation features to be included in subsequent versions, that will be analyzed following the same pipeline and be permanently stored in public repositories. We believe that MATEdb will accelerate research on animal phylogenomics while saving thousands of hours of computational work in a plea for open and collaborative science.

## Introduction

### Phylotranscriptomics and the quest for resolving the Animal Tree of Life

With the advent of high throughput sequencing techniques, genomic approaches to phylogenetics, and more specifically analyses based on transcriptomic data (commonly referred to as phylotranscriptomics), are quickly replacing single or few-gene approaches, enabling an in-depth phylogenetic interrogation of multiple lineages at an unprecedented level in a reliable manner (Cheon, Zhang, and Park 2020). Since the cost of sequencing and assembling transcriptomes is still much lower compared to genomes, this source of genomic data has been extensively exploited with the goal of accessing thousands of molecular markers that can be leveraged to explore phylogenetic interrelationships. For example, the inference of phylogenetic relationships based on a handful of genes has historically failed to provide enough resolution to illuminate animal interrelationships at both deep and shallow levels in many metazoan lineages. Conversely, phylotranscriptomic approaches have been successfully used to explore phylogenetic relationships within phyla in multiple animal lineages such as arthropods (Schwentner et al. 2017; Fernández et al. 2018; Lozano-Fernandez et al. 2019), annelids (Novo et al. 2016; Erséus et al. 2020), molluscs (Kocot et al. 2011; Zapata et al. 2014), echinoderms (Mongiardino Koch et al. 2018) or nematodes (Smythe, Holovachov, and Kocot 2019), among others. In addition, these approaches have also served to advance our understanding of the deepest relationships of the Animal Tree of Life (Laumer et al. 2019).

### Leveraging transcriptomes to interrogate gene repertoire evolution across animal lineages

The plethora of transcriptomic datasets available in public databases can potentially be harnessed not only to understand phylogenetic interrelationships but also to investigate the dynamics of gene repertoire evolution. However, its immediate use would require effective data mining and integration. More specifically, since most genes are expressed basally in most tissues (Ramsköld et al. 2009; Gu et al. 2021), transcriptomic data from a species can be used as a proxy of its proteome. The combination and joint analysis of data from several species allows the interrogation of the dynamics of gene family evolution at macroevolutionary scales. This approach has been followed to investigate evolutionary dynamics in complex gene families in clades such as land plants (Geng et al. 2021), insects (Thoma et al. 2019) or molluscs (De Oliveira et al. 2016), and has also been leveraged to investigate macroevolutionary patterns of gene gain and loss across animal phyla (De Oliveira et al. 2016; Fernández and Gabaldón 2020).

Unique among related databases, we present here Metazoan Assemblies from Transcriptomic Ensembles (MATEdb v1), a continuously updated and curated database of hundreds of high-quality transcriptome assemblies from different animal phyla. The database included both retrieved raw transcriptomic data from public databases and datasets generated by us. We performed a *de novo* transcriptome assembly and after a quality filtering process, we evaluated the integrity score of the gene content against the scores of the BUSCO database. We complement the repository with high-quality genome gene annotations, which are often not readily available from sequencing projects. We provide orthology-based gene annotations for all datasets as well. The main differences between MATEdb and other databases (e.g., MolluscDB, Liu et al. 2021) are as follows: (i) the datasets included in MATEdb are all high quality (i.e. high BUSCO completeness). This is key to minimize biases during downstream analyses such as orthology inference or gene repertoire evolution studies; (ii) all datasets in MATEdb have been analyzed with the exact same pipeline. This is to our belief one of the most valuable features of our resource, since different versions of the same software (e.g. Trinity) change substantially (e.g. in our experience the number of ‘genes’ inferred with Trinity may vary up to one order of magnitude depending on the version); (iii) MATEdb pays more attention to lineage representation than to the total number of datasets included, i.e. we prioritize the inclusion of the main lineages within each phyla over the number of species included. We believe that MATEdb is a valuable resource that will promote and accelerate research on animal evolutionary studies while saving thousands of hours of computational work in a call for open and collaborative science.

## Methods

### Taxonomic coverage

Our goal is to provide a comprehensive data repository of high quality genomic and transcriptomic spanning all animal phyla with emphasis in a good representation of the main lineages within each phylum. We intend this manuscript to be the core of the database and the general description, and aim at updating it with new versions expanding to other animal phyla in the near future. The first version of MATEdb (v1) includes arthropods and molluscs. Taxon sampling is represented in Figure 1 and Table S1 (these files will be updated for the subsequent versions of the database). Taxon sampling was based on dataset availability in public repositories while maximizing taxonomic representation at the order and family level over closely related species. The wet lab protocol for newly generated data is described in Suppl. File S1 (in folder ‘Protocols’).

**Figure 1.**
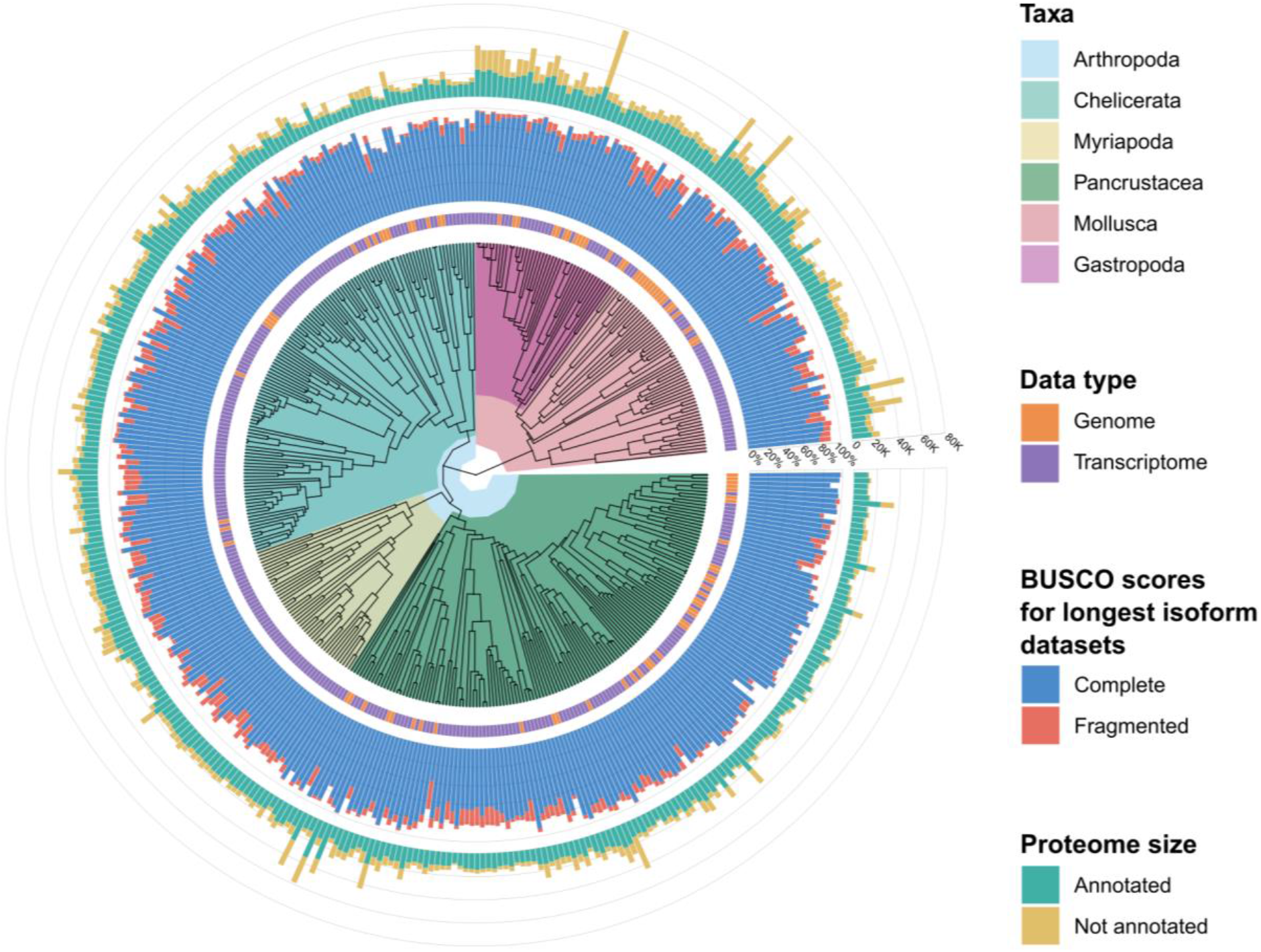
Taxonomic representation of taxa included in MATEdb v1. Taxonomic rank (e.g., phylum, class, etc.), data type (i.e., genome or transcriptome), BUSCO score for the longest isoform datasets and proteome size is shown in the outer rings.

### Analytical pipeline

The analytical pipeline followed is depicted in Figure 2. In brief, the main steps were as follows: (1) raw data were downloaded from public repositories (most of them from the Sequence Read Archive (SRA) in NCBI); (2) adapters and low-quality reads were trimmed; (3) filtered raw reads were de novo assembled; (4) assembly completeness was explored based on the percentage of metazoan single-copy genes recovered; (5) Open Reading Frames (ORFs) were inferred from the assemblies; (6) ORFs were decontaminated (e.g., filtering out non-metazoan ORFs); (7) the longest isoform per gene (sensu Trinity) was parsed, which can be directly used as the input for phylogenomic studies. Further details about the software versions and parameters implemented throughout the pipeline are shown in Figure 2 (see also Table S2).

**Figure 2.**
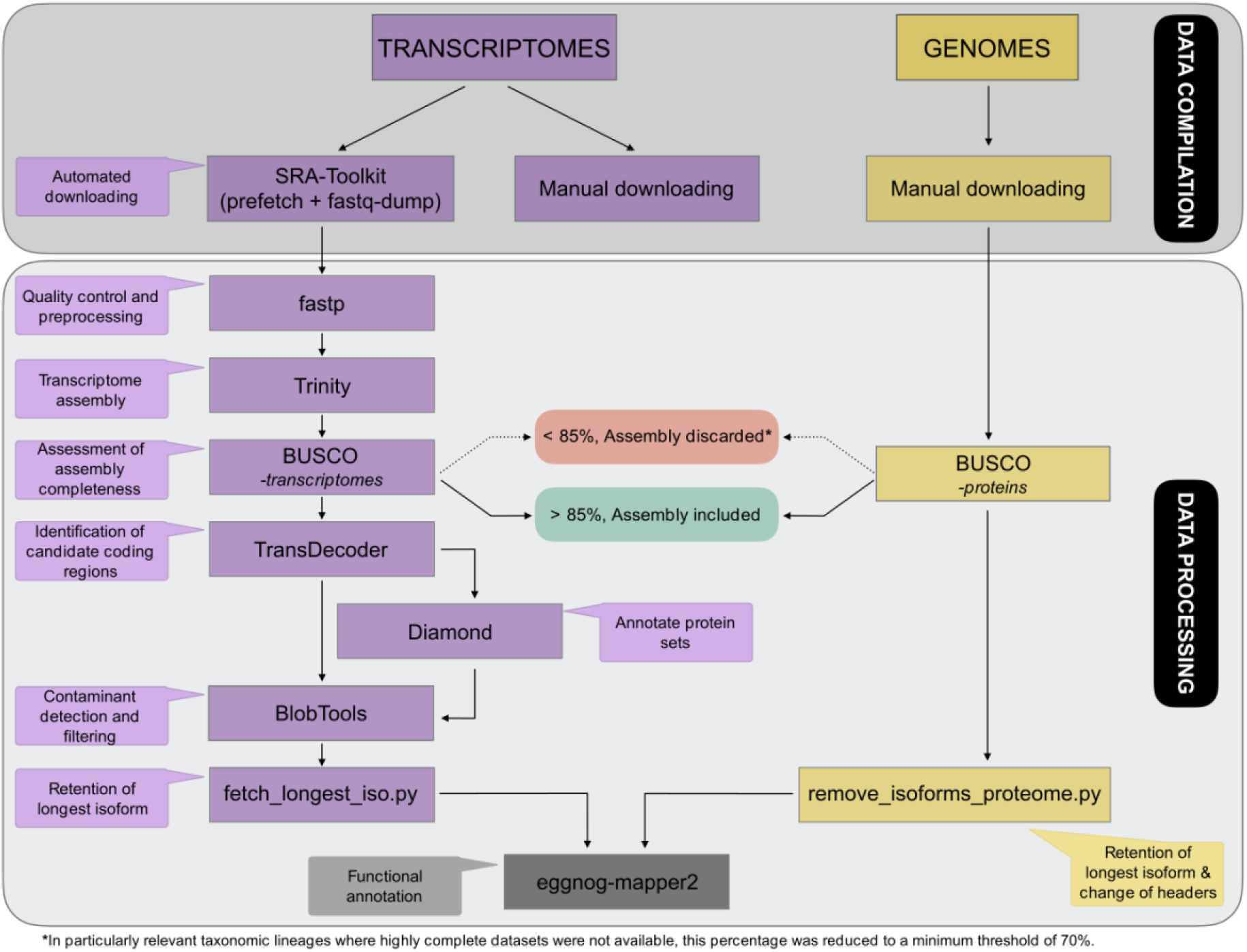
Pipeline followed to generate the MATEdb database. All steps are discussed in detail below.

### Compilation of transcriptomic data ensembles

RNA-seq raw data for the selected samples (accession numbers are available as Supplementary Material, Table S1) were retrieved from SRA (Leinonen et al. 2011) by using the fastq-dump program of the SRA Toolkit v.2.10.7 (http://ncbi.github.io/sra-tools/). Paired-end sequences were preferably chosen over single reads when possible. Moreover, raw data published in MolluscDB (Liu et al. 2021 and Caurcel et al. 2021) repositories were manually downloaded.

For the projects in which transcriptome data sequencing was not extracted from the whole specimen but on specific tissues, a pool was created to combine all raw data referred to the same organism and the same project. Each accession was independently downloaded, although they were considered a unique data set for the rest of the analyses.

### Compilation of genomic data

Coding DNA Sequences (CDS) and proteome files including all predicted proteins within genome assembly were downloaded from the different data repositories referenced in Supplementary Material, Table S1.

### Transcriptome assembly processing

Downloaded sequenced reads were checked for quality using package fastp v.0.20.0 (https://github.com/OpenGene/fastp; Chen *et al*. 2018) by trimming adapters and primers, filtering reads with phred quality <Q5, and filtering reads with N base number >5. De novo assembly of clean data was constructed using Trinity v2.11 (Grabherr et al. 2011) with default parameters. After Trinity execution, all headers in resultant assemblies were normalized by adding custom unique code plus gene and isoform identifiers for each sequence (see Supplementary Material, Table S2). For the cases in which transcriptomes consisted of a pool of libraries belonging to a single specimen, each of the datasets was trimmed independently and then merged together during the assembly process with Trinity.

We assessed the quality of assembled transcriptomes using completeness scores of BUSCO v4.1.4 (Manni et al. 2021) with the metazoa_db10 (for molluscs) and the arthropoda_db10 (for arthropods). BUSCO software employs sets of benchmarking universal single-copy orthologs from OrthoDB (www.orthodb.org; accessed on 10 February 2021) to provide quantitative measures of the completeness of genomic assemblies in terms of expected gene content.

Transcriptome assemblies with a gene completeness percentage (considering the sum of both complete and fragmented genes) higher than 85% were retained. As an exception to this, there are some transcriptome assemblies which have been included despite their low BUSCO score due to their taxonomic relevance (e.g., they were the only representatives of their lineage, such as in the case of Remipedia). In any case, the lowest threshold for BUSCO scores allowed in MATEdb was ca. 70%.

Candidate coding regions were predicted on assemblies, using TransDecoder v5.5.0 (https://github.com/TransDecoder/TransDecoder) in two different steps aiming to improve the reliability of the prediction. First, all ORFs with a minimum length of 100 amino acids were extracted with the TransDecoder.LongOrfs module. Then, 25% of the total number of previously predicted ORF was computed and used as input to train the Markov Model behind the TransDecoder.Predict module, which resulted in a more accurate output than the one done with the default settings. For further details see Supplementary Material, Table S2.

Contaminant sequences present in the transcriptome assemblies were filtered out based on BlobTools2 analysis (Challis et al. 2020). First, assemblies were subjected to a Diamond (Buchfink, Xie, and Huson 2015) BLASTP search (--sensitive --max-target-seqs 1 --evalue 1e-10) against the NCBI non-redundant (NCBI NR) protein sequence database (http://www.ncbi.nlm.nih.gov/). Contigs that were classified as bacterial, plant, or fungal sequences were removed from the assembly. All commands needed during the decontamination process with BlobTools were applied through the execution of several custom scripts (“blobtools.sh” and “extract_phyla_for_blobtools.py”), included in the Github repository.

Finally, the longest ORFs of each transcript were retained as final candidate coding regions for further analyses. This step was performed by applying a modified version of the python script “choose_longest_iso.py” from (Fernández et al. 2014) to adapt to the new Trinity headers, also included in the Github repository (“fetch_longest_iso.py”).

#### Genome data processing

We downloaded the predicted proteome for published genomes mostly from the NCBI genome library (https://www.ncbi.nlm.nih.gov/genome/). Then, gene completeness was assessed using BUSCO in protein mode. As it was established for transcriptome assemblies, only those proteome assemblies with BUSCO completeness score higher than 85% (complete plus fragmented percentages). In addition, we carried out a filtering step to keep the longest isoform of each gene using a custom script similar to that used with transcriptomes filtering (“remove_isoforms_proteome.sh”). This script takes the annotation file as a reference to create the new Trinity-like headers and finally returns the filtered proteome file. For further details on scripts, please check the Github repository.

### Functional annotation of gene repertoire

The longest isoform gene list for each dataset was annotated with eggNOG-mapper v2 (Cantalapiedra et al. 2021). This permits a higher precision than traditional homology searches (i.e. BLAST searches), as it avoids transferring annotations from close paralogs (duplicate genes with a higher chance of being involved in functional divergence).

## Database availability

### Scripts and commands

The scripts and commands used for every step and the supplementary Tables S1 and S2 are publicly available in the following repository: https://github.com/MetazoaPhylogenomicsLab/MATEdb

### Files deposited in the repository

The data repository is composed of (1) a de novo transcriptome assembly, (2) its candidate coding regions within transcripts (both at the level of nucleotide and amino acid sequences), (3) the coding regions filtered using their contamination profile (i.e., only metazoan content), (4) the longest isoform of the amino acid candidate coding regions, (5) the gene content completeness score as assessed against the BUSCO database, and (6) an orthology-based gene annotation. We complement the repository with genome annotations from high-quality genomes that are often not straightforward to obtain from the sequencing projects (in the case of genomes, only files (4), (5) and (6) are provided in MATEdb).

### Software availability

We provide a Docker container for easy deployment of the tools used to generate the files in the database with the appropriate software versions along with their dependencies (https://hub.docker.com/repository/docker/vargaschavezc/matedb). The software included is the following: SRA Toolkit version 2.10.7, fastp version 0.20.1, Trinity version 2.11.0, BUSCO version 4.1.4, TransDecoder version 5.5.0, Diamond 2.0.8, BlobTools 2.3.3 and eggNOG-mapper 2.1.6.

## Author contribution

This database is the result of the collaborative effort of lab members from the Metazoa Phylogenomics Lab to offer the scientific community the possibility to reuse some of the data generated for their individual projects. VT, CSG, JL-F, GIMR, PBG, LA and KE contributed assemblies to the data repository. Author order is based on the overall contribution to the database, with female lab members listed first in case of similar contributions. GIMR created the pipeline custom scripts for the genome data analyses and designed the MATEdb logo. CVC and RF contributed to the creation and management of the database. CVC created and curated the Github repository and prepared the Docker container. RF provided resources and wrote the first version of the manuscript. All authors contributed to the writing and approved the manuscript.

## Acknowledgements

Preprint version 4 of this article has been peer-reviewed and recommended by Peer Community In Genomics (https://doi.org/10.24072/pci.genomics.100022). We thank Centro de Supercomputación de Galicia (CESGA) for access to computer resources, and particularly Pablo Rey for his kind assistance and guidance.

## Data, scripts and codes availability

Data, scripts and code are available online: https://github.com/MetazoaPhylogenomicsLab/MATEdb The docker container is available online: https://hub.docker.com/repository/docker/vargaschavezc/matedb

## Supplementary material

Supplementary material are available online: https://github.com/MetazoaPhylogenomicsLab/MATEdb

## Conflict of interest disclosure

The authors declare that they comply with the PCI rule of having no financial conflicts of interest in relation to the content of the article. RF is a recommender and a member of the managing board of PCI Genomics.

## Funding

PBG was supported by an FPI grant (grant agreement no. BES-2017-081050) financed by MCIN/AEI /10.13039/501100011033 and by European Social Fund (ESF) ‘Investing in your Future’ and the Systematics Research Fund 2020 (Linnean Society of London and the Systematics Association). GIMR acknowledges the support of Secretaria d’Universitats i Recerca del Departament d’Empresa i Coneixement de la Generalitat de Catalunya and ESF Investing in your future (grant 2021 FI_B 00476). LA and JL-F acknowledge funding from a Juan de la Cierva-Formación and Juan de la Cierva-Invorporación (grant agreements FJC2019-042184-I and IJC2018-035237-I respectively, funded by MCIN/AEI/10.13039/501100011033). RF acknowledges support from the following sources of funding: Ramón y Cajal fellowship (grant agreement no. RYC2017-22492 funded by MCIN/AEI /10.13039/501100011033 and ESF ‘Investing in your future’), the Agencia Estatal de Investigación (project PID2019-108824GA-I00 funded by MCIN/AEI/10.13039/501100011033) and the European Research Council (this project has received funding from the European Research Council (ERC) under the European’s Union’s Horizon 2020 research and innovation programme (grant agreement no. 948281)).

## Notes

https://github.com/MetazoaPhylogenomicsLab/MATEdb

### Competing Interest Statement

The authors have declared no competing interest.

### Summary of Updates

Version 4 reviewed and recommended by PCI Genomics

